# iELVis: An open source MATLAB toolbox for localizing and visualizing human intracranial electrode data

**DOI:** 10.1101/069179

**Authors:** David M. Groppe, Stephan Bickel, Andrew R. Dykstra, Xiuyuan Wang, Pierre Mégevand, Manuel R. Mercier, Fred A. Lado, Ashesh D. Mehta, Christopher J. Honey

**Affiliations:** Dpartment of Psychology, University of Toronto, Toronto, ON M5S SG3, Canada; Department of Neurosurgery, Hofstra Northwell School of Medicine, and Feinstein Institute for Medical Research, Manhasset, NY 11030, USA; Department of Neurology, Montefiore Medical Center, Bronx, NY 10467, USA; Department of Neurology, Stanford University, Stanford, CA 94305, USA; Department of Neurology, Ruprecht-Karls-Universität Heidelberg, 69120 Heidelberg, Germany; Department of Neurology, New York University School of Medicine, New York, NY 10016, USA; Department of Radiology, New York University School of Medicine, New York, NY 10016, USA; Division of Neurology, Department of Clinical Neuroscience, Hôpitaux Universitaires de Genève, 1211 Geneva, Switzerland; Centre de Recherche Cerveau et Cognition (CerCo), CNRS, UMR5549, Pavillon Baudot CHU Purpan, BP 25202, 31052 Toulouse Cedex, France; Université Paul Sabatier, Toulouse, France; Department of Neuroscience, Albert Einstein College of Medicine, Bronx, NY 10461, USA

**Author notes:** Correspondence and requests for materials should be addressed to: David M. Groppe, Ph.D. Dept. of Psychology University of Toronto 100 St. George St. Toronto, ON M5S SG3 Canada.

## Abstract

**Background:** Intracranial electrical recordings (iEEG) and brain stimulation (iEBS) are invaluable human neuroscience methodologies. However, the value of such data is often unrealized as many laboratories lack tools for localizing electrodes relative to anatomy. To remedy this, we have developed a MATLAB toolbox for intracranial electrode localization and visualization, iELVis.

**New Method:** iELVis uses existing tools (BioImage Suite, FSL, and FreeSurfer) for preimplant magnetic resonance imaging (MRI) segmentation, neuroimaging coregistration, and manual identification of electrodes in postimplant neuroimaging.

Subsequently, iELVis implements methods for correcting electrode locations for postimplant brain shift with millimeter-scale accuracy and provides interactive visualization on 3D surfaces or in 2D slices with optional functional neuroimaging overlays. iELVis also localizes electrodes relative to FreeSurfer-based atlases and can combine data across subjects via the FreeSurfer average brain.

**Results:** It takes 30-60 minutes of user time and 12-24 hours of computer time to localize and visualize electrodes from one brain. We demonstrate iELVis’s functionality by showing that three methods for mapping primary hand somatosensory cortex (iEEG, iEBS, and functional MRI) provide highly concordant results.

**Comparison with Existing Methods:** iELVis is the first public software for electrode localization that corrects for brain shift, maps electrodes to an average brain, and supports neuroimaging overlays. Moreover, its interactive visualizations are powerful and its tutorial material is extensive.

**Conclusions:** iELVis promises to speed the progress and enhance the robustness of intracranial electrode research. The software and extensive tutorial materials are freely available as part of the EpiSurg software project: https://github.com/episurg/episurg

## 1 INTRODUCTION

Since the early 20^th^ century, the intracranial electroencephalogram (iEEG) and intracranial electrical brain stimulation (iEBS) have been invaluable tools for mapping pathological and functional brain regions in patients being evaluated for resective brain surgery (Penfield & Jasper, 1954). In addition to their clinical utility, these invasive measures have provided a unique window on human brain function for both clinical and basic research.

There has been a dramatic increase of interest in iEEG and iEBS research over the past decade, for several reasons. Firstly, iEEG has proven to be a fine-grained measure of mean local firing rates (Crone, Korzeniewska, & Franaszczuk, 2011; Miller, Sorensen, Ojemann, & Nijs, 2009) as well as local synaptic potentials (e.g., Golumbic et al., 2013). Secondly, there is mounting evidence that iEBS can improve (Suthana et al., 2012) or manipulate (Parvizi et al., 2012; Mégevand et al., 2014) brain function with great precision. Finally, a growing number of public databases of iEEG data (e.g., www.ieeg.org, www.epilepsiae.eu) are increasing access to these rare data.

Despite the growing popularity iEEG and iEBS research, many researchers encounter technical challenges when attempting to analyze these data. These include: [1] precisely identifying the anatomical location of electrodes, [2] correcting for postimplant brain deformities (i.e., “brain shift”) so that postimplant data can be coregistered to preimplant neuroimaging, [3] effectively visualizing electrode data in a way that communicates their locations relative to other neural measures (e.g., magnetic resonance imaging, MRI, and functional magnetic resonance imaging, fMRI), and [4] mapping idiosyncratic electrode montages into a common space for multi-patient analyses. Although some excellent public software has been produced that solves some of these issues ((Table 1), there is not yet a well-developed, scriptable, public software package for solving all of them. Consequently, iEEG and iEBS research groups currently develop lab-specific solutions from existing packages and their own custom code. This inefficient and error-prone solution to localization and visualization makes it difficult to compare findings across research groups.

**Table 1.**
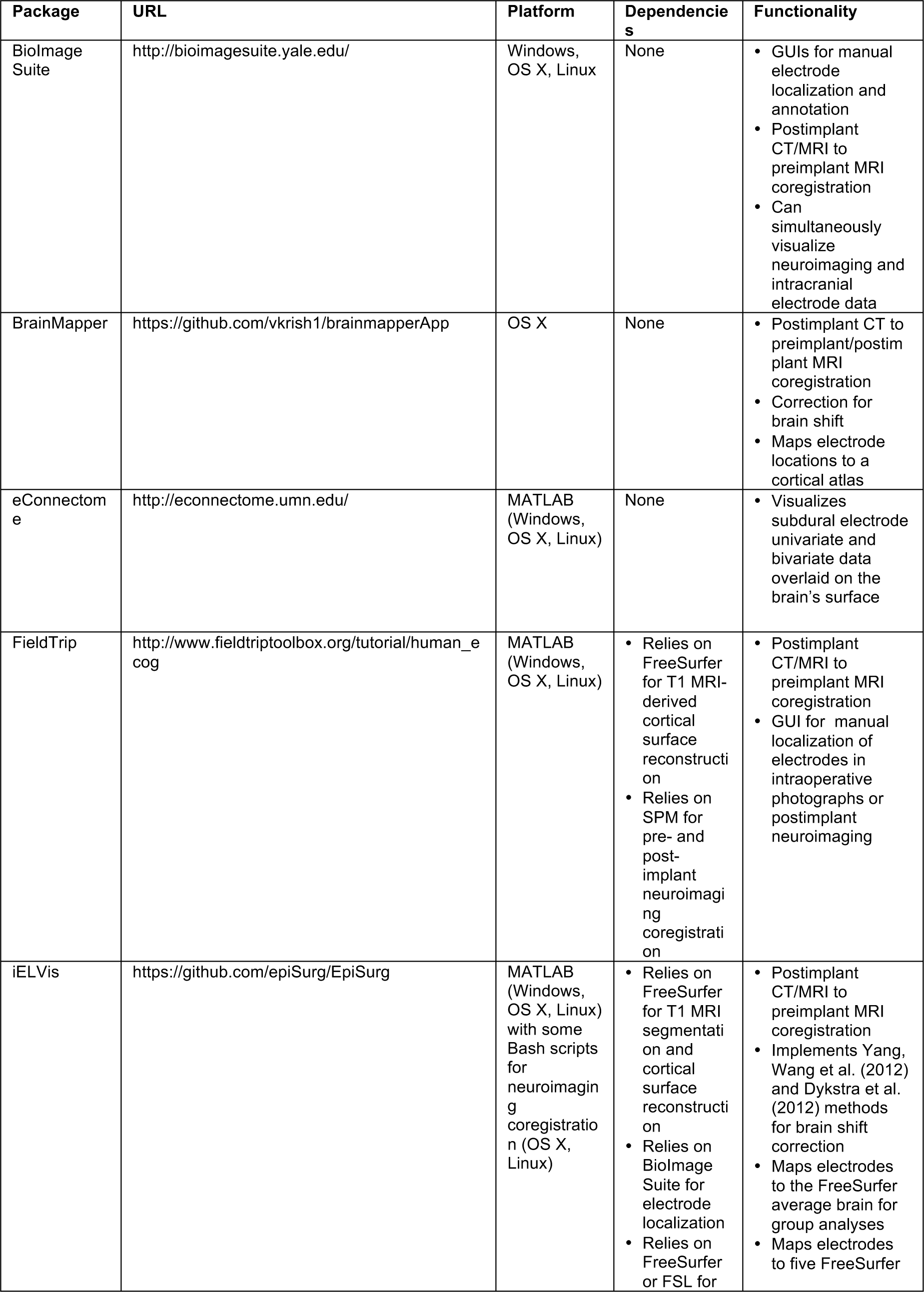
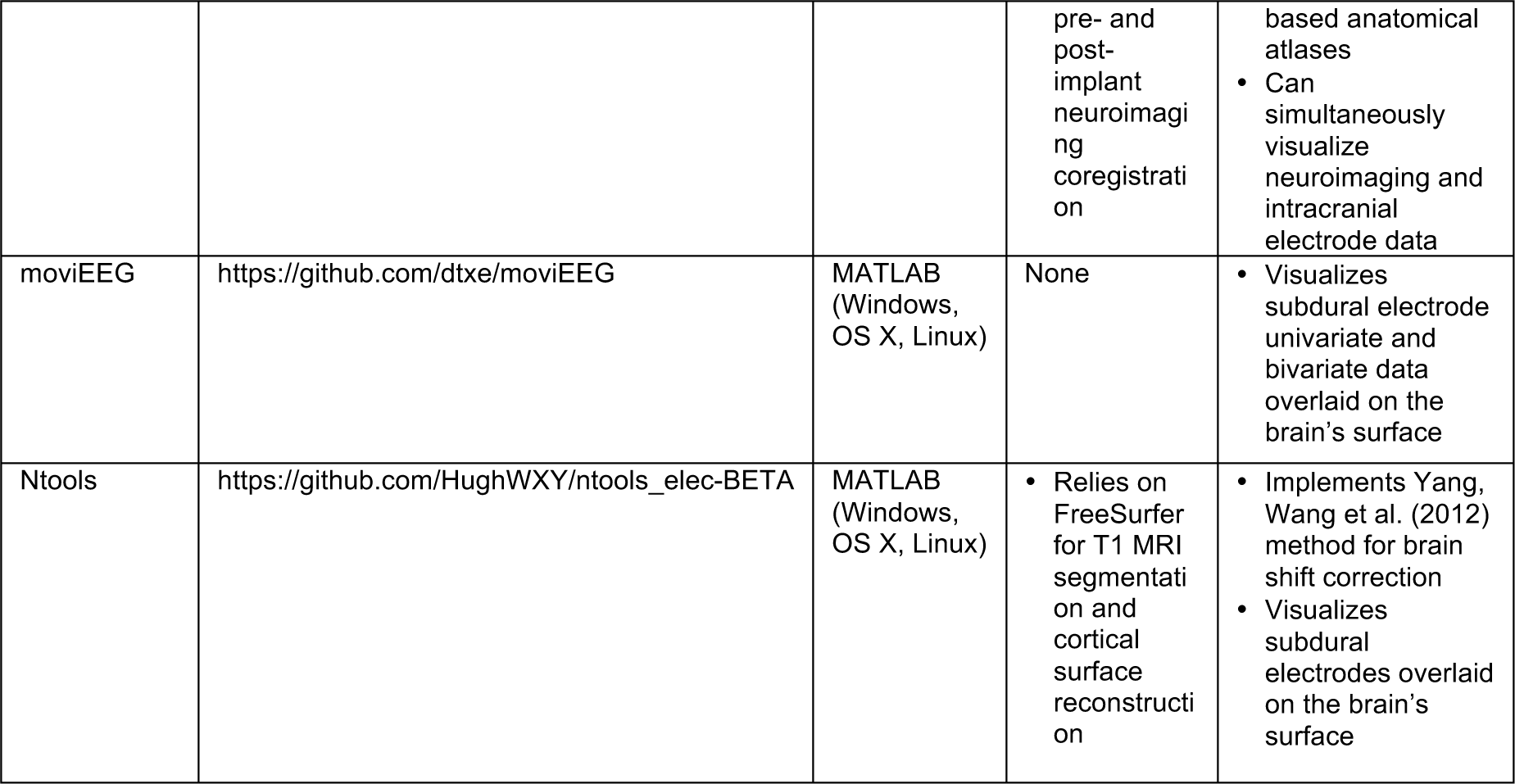
Some existing software packages that support intracranial electrode localization or visualization.

To remedy this problem, we have pooled the resources of five different iEEG/iEBS research groups to develop a freely-available, open source software toolbox and processing pipeline that can assist researchers in localizing electrodes, correcting for brain shift, overlaying electrode locations with neuroimaging data, and visualizing their locations relative to individual anatomy and group-level templates. The toolbox, called iELVis (Intracranial ELectrode VISualization), consists of MATLAB functions with a handful of Bash scripts for neuroimaging coregistration and overlay creation. iELVis relies on FreeSurfer (www.freesurfer.net) for MRI segmentation and for mapping to the FreeSurfer average brain for group analyses. In addition, iELVis relies on BioImage Suite (www.bioimagesuite.org; Papademetris, Jackowski, & Rajeevan, 2011) for manually tagging electrode locations in postimplant CT or MRI scans. and FSL or FreeSurfer for coregistering pre-and post-implant neuroimaging.

The key contributions of iELVis are that it:

1. implements a set of standard tools for electrode localization/visualization, using MATLAB code supported by thorough tutorials and documentation
2. establishes a pipeline and toolset to standardize localization across multiple laboratories
3. implements new quality control metrics to help identify erroneous localizations
4. produces interactive, multi-view representations of the brain that can simultaneously represent multiple data modalities (e.g., iEEG, iEBS, & fMRI)

This paper describes the main features and workflow of iELVis. More extensive documentation and tutorial materials are available on the toolbox wiki, http://episurg.pbworks.com. This includes two example datasets and example code that illustrate how to use all the functionalities described here. The toolbox can be downloaded from https://github.com/episurg/EpiSurg.

## 2 ELECTRODE LOCALIZATION

The first step in electrode localization is to preprocess and automatically segment a patient’s preimplant T1 MRI using FreeSurfer. The preprocessing aligns the MRI to a standard FreeSurfer coordinate space that is used for all iELVis processing and visualization. The segmentation assigns each brain voxel to one of 46 regions such as the hippocampus, amygdala, cerebral cortex, or white matter (Figure 1: A left). The segmentation also estimates the pial surface for each cerebral hemisphere and smooths over the sulci in the pial surface to derive a proxy for the leptomeningeal surface^1^ (Figure 1: A middle & right). The leptomeningeal surface (Schaer et al., 2008) is created because it is useful for identifying subdural electrode locations since subdural electrodes traverse sulci. Finally, the individual’s brain is mapped to the FreeSurfer average brain (see subsequent section). This procedure requires up to 24 hours on a conventional workstation. Manual intervention is rarely necessary. The exceptions are patients with gross brain abnormalities such as tumors and lobectomies for whom the automatic segmentation may fail around the abnormal regions, and the medial wall of the anterior medial temporal lobe, which tends to be underestimated.

**Figure 1:**
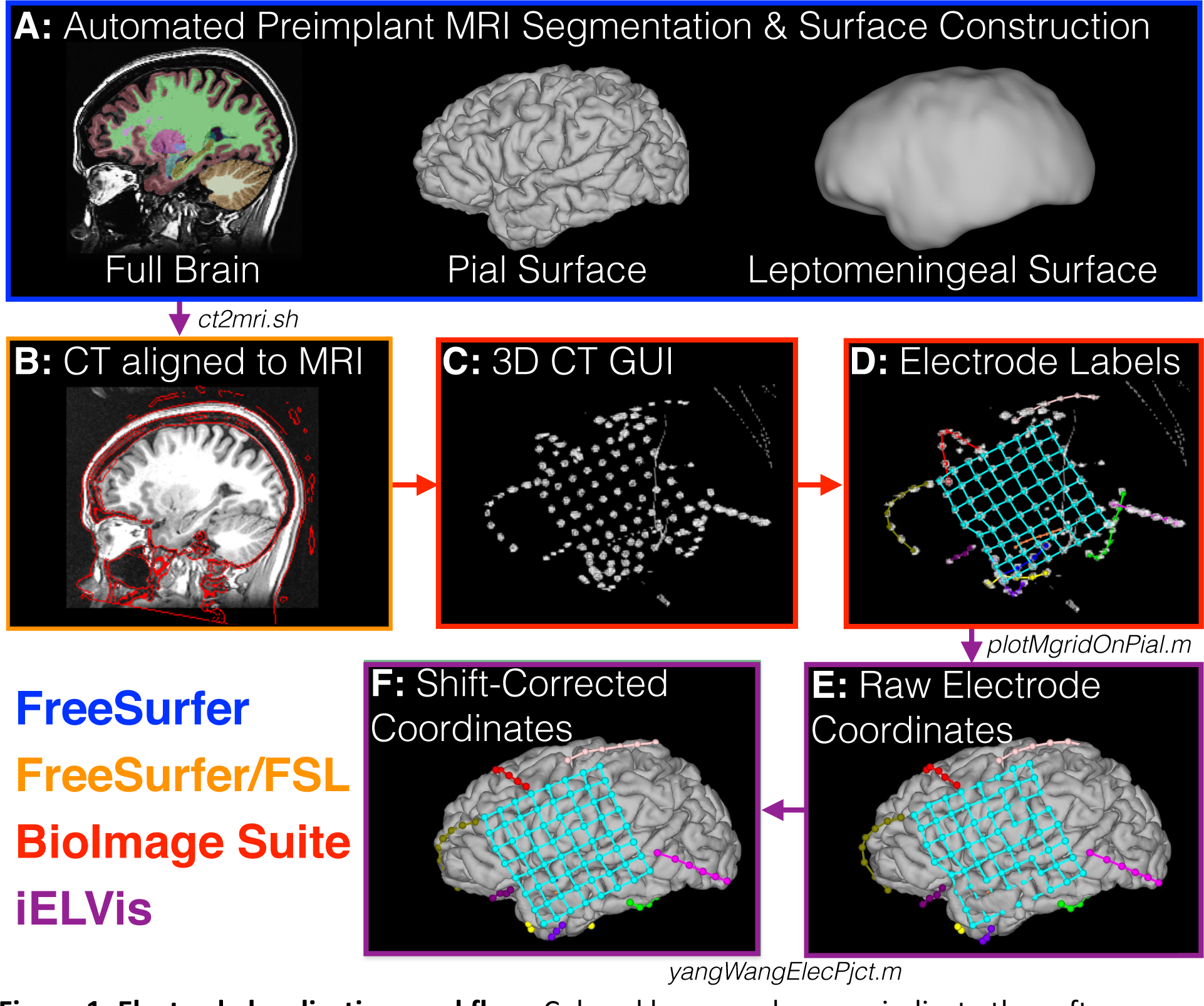
Electrode localization workflow. Colored boxes and arrows indicate the software package used for each workflow step (FreeSurfer, FSL, BioImage Suite, or iELVis). Italicized words between boxes indicate iELVis functions used to implement that processing step. **[A]** FreeSurfer automatically segments various anatomical regions from a preimplant T1 MRI. This produces a 3D labeling of brain structures such as white matter, hippocampus, and neocortical grey matter which are represented with different colors. In addition, FreeSurfer constructs pial and leptomeningeal surfaces. **[B]** A postimplant CT scan is rigidly coregistered to the preimplant T1 MRI. Edges of the CT scan are overlaid as red lines on the T1 MRI. **[C]** Electrodes are clearly apparent in the CT scan, visualized in 3D via a BioImage Suite graphical user interface (GUI). **[D]** Electrode locations are manually labeled using BioImage Suite. Each strip, grid, or shaft of electrodes is represented with a unique color. **[E]** Electrode locations on the FreeSurfer reconstructed pial surface visualized with iELVis functions in MATLAB. Some cyan grid electrodes appear to lie inside the temporal lobe, due to postimplant brain compression (“brain shift”). **[F]** Brain shift-corrected electrode locations on the FreeSurfer pial surface visualized with iELVis functions in MATLAB. Note that all data are shown in the standard FreeSurfer coordinate space for the subject.

Once FreeSurfer has completed processing the MRI, a postimplant computerized tomography (CT) or MRI scan is rigidly aligned to the preimplant MRI via an affine transform with six degrees of freedom (Figure 1: B). iELVis provides Bash scripts for performing this coregistration using the *flirt* tool (Jenkinson & Smith, 2001; Jenkinson, Bannister, Brady, & Smith, 2002) from the Oxford Centre for Functional MRI of the Brain Software Library (FSL: www.fmrib.ox.ac.uk/fsl) or using FreeSurfer’s *bbregister* (Greve & Fischl, 2009). While *flirt* attempts to align the entire volume, *bbregister* aligns image boundaries. Another practical difference is that the results of *bbregister* are easy to manually edit using FreeSurfer’s *tkregister2* graphical user interface (GUI). Accuracy of the CT-MRI coregistration can be readily visually verified by the alignment of the skull in both volumes. MRI-MRI coregistration accuracy is even easier to visually confirm. In our testing, both *flirt* and *bbregister* typically give very similar results with no need for manual intervention. However, when one method fails, the other will almost always succeed.

After the postimplant scan has been aligned to the preimplant MRI, the postimplant scan (in the preimplant MRI space) is imported into BioImage Suite’s Electrode Editor GUI to manually identify electrode locations (Figure 1: C-D). Depending on the number of electrodes, a user can typically tag all electrodes in 30 to 60 minutes. Next, electrode coordinates are saved to a BioImage Suite *mgrid* file which can be imported into MATLAB and visualized over the pial surface using iELVis functions (Figure 1: E). Other neuroimaging interfaces (e.g., FreeSurfer’s *tkmedit*) could be used in lieu of BioImage Suite for identifying electrode coordinates if users record electrode locations in a text file compatible with iELVis conventions.

Finally, the subdural electrodes are projected out to the leptomeningeal (i.e., the smoothed pial) surface to correct for brain shift (Figure 1: F). Brain shift is caused by factors such as loss of cerebrospinal fluid, swelling, and the displacement of the brain by electrodes (Hastreiter et al., 2004). This deformity is typically most severe near a craniotomy and can be more than 1 cm (Dalal et al., 2008; Hill, Smith, & Simmons, 2000). In contrast, implants requiring only burr holes typically produce minimal brain shift (Sweet, Hdeib, Sloan, & Miller, 2013). iELVis includes two algorithms for brain shift correction. The first of these, devised by Dykstra, Chan and colleagues (Dykstra et al., 2011), projects each subdural electrode to the leptomeningeal surface using an iterative optimization algorithm that attempts to minimize the change in each electrode’s location and the distance with its one to five closest neighbors. Based on comparisons with intraoperative photographs of electrode locations in five patients, the Dykstra algorithm appears to localize electrodes under the craniotomy with median (interquartile range) error of 3 (2.39) mm or less (ibid.). Since electrodes near the craniotomy are typically most affected by brain shift, accuracy is likely even better for strips of electrodes that are inserted far from or without craniotomies. The second algorithm was created by Yang, Wang and colleagues (Yang et al., 2012) and projects grids of electrodes to the leptomeningeal surface via an inverse gnomonic projection.

More specifically, the algorithm approximates the leptomeningeal surface under each grid as part of a larger sphere and iteratively adjusts the projection of the grid plane onto the sphere to minimize the difference between the projected and known electrode geometry. Subdural strips of electrodes are simply assigned to the location of the nearest point on the leptomeningeal surface. When the results of this algorithm were compared with electrode locations in intraoperative photographs in eight patients, the median (interquartile range) error of the Yang-Wang method was 0.74 (0.75) mm for sub-craniotomy electrodes (Yang et al., 2012). Once the subdural electrodes have been corrected for brain shift, several diagnostic plots are produced to help identify potential errors (Figure 2).

**Figure 2:**
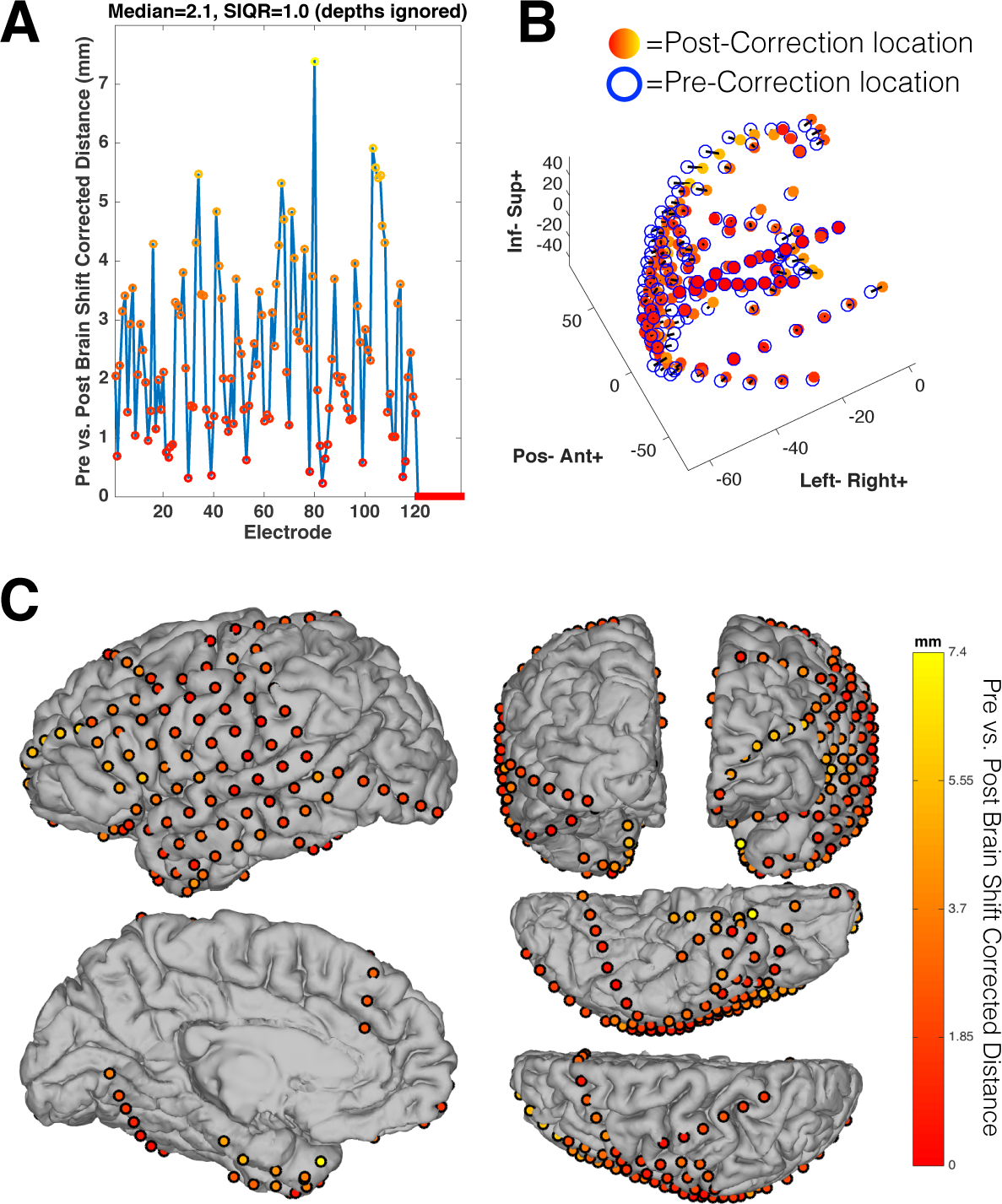
Correcting for brain shift using iELVis. **[A]** The distance between electrode locations before and after brain shift correction. Subdural electrodes (circles) are corrected for brain shift, while depth electrodes (squares) are not; the correction distance for depth electrodes is therefore zero. Color scale corresponds to the plot in C. **[B]** Electrode locations pre-and post-correction for brain shift; lines join pre-and post-correction locations of each electrode. Post-correction electrodes use the same color scale as the plot in C. [**C**] Electrode locations overlaid on the subject’s pial surface, after brain shift correction, color coded to reflect the correction distances in A. In all figures, clicking on electrode symbols will produce the corresponding electrode’s name. The three dimensional figures can be interactively rotated.

Note that neither algorithm corrects the location of penetrating depth electrodes for brain shift. Since depth electrodes typically target medial structures (e.g. the hippocampus) that are minimally affected by brain shift, this is likely not a serious shortcoming. Moreover, depth-only implants (i.e., stereotactic EEG) generally produce minor brain shift as a craniotomy is not performed (Gonzalez-Martinez et al., 2014; Sweet, Hdeib, Sloan, & Miller, 2013).

## 3 MAPPING ELECTRODES TO AN AVERAGE BRAIN

iELVis supports mapping electrodes to the FreeSurfer average brain for combining data across patients (i.e., “group analyses”). FreeSurfer maps individual brains to its average brain by aligning their gyrification patterns. This mapping thus accurately preserves the gyral location of subdural electrodes at the cost of greatly distorting some inter-electrode distances (Figure 3).

**Figure 3:**
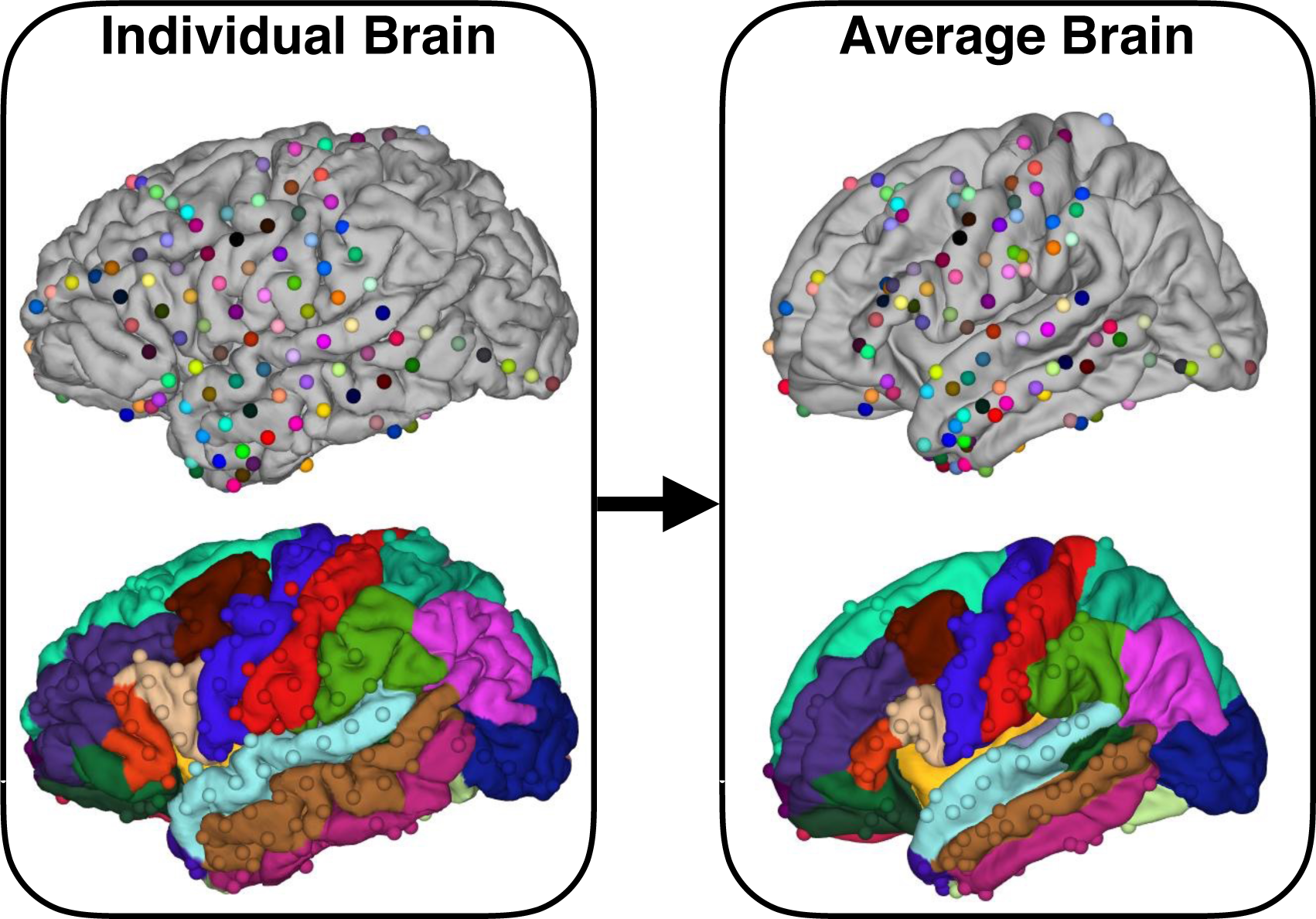
Mapping from individual anatomy to the FreeSurfer average brain preserves gyral location. **[Left-Top]** Electrodes on an individual’s pial surface are colored uniquely. **[Left-Bottom]** The individual brain is colored according to cortical areas as defined by FreeSurfer’s Desikan-Killiany atlas. Electrodes are colored to match that of the closest brain region. **[Right]** Analogous electrode locations on the FreeSurfer average brain with the same color codes used for the individual brain. Mapping to the average brain preserves the gyral location of each electrode at the cost of distorting electrode geometry. Note that the FreeSurfer average brain has exaggerated sulci because the gyri-aligned inter-subject averaging that produced it smooths gyrification patterns

As described in Dykstra et al. (2012), iELVis maps subdural electrode locations from a patient’s brain to the average brain by first assigning each electrode to the closest point on the patient’s pial surface (after the leptomeningeal surface-based brain shift correction described above). The pial surface is warped to a sphere which FreeSurfer has gyrally aligned to a spherical version of its average brain’s pial surface (Fischl et al., 1999). Each point on the patient’s spherical surface is assigned to the nearest neighbor on the average brain spherical surface, which has a one-to-one vertex correspondence to the average brain pial surface.

By default, penetrating depth electrodes are mapped to the FreeSurfer average brain via an affine transformation to MNI305 space, which is compatible with that of the FreeSurfer average brain. However, penetrating electrodes within deep structures such as the amygdala and hippocampus might be best grouped across patients by using volumetric anatomical atlases (see following section). Depth electrode contacts that lie in cerebral grey matter can be mapped to the average brain via surface based mapping if users manually edit the iELVis electrode location files to indicate that the contact is a subdural electrode instead of a depth electrode.

## 4 CATEGORIZING ELECTRODES USING ANATOMICAL ATLASES

In addition to mapping individual brains to standard spaces, it is often useful to map them to standard anatomical atlases. Anatomical atlases can be used to define regions of interest to select subsets of electrodes for analysis or to combine data across patients for group analyses (e.g., Groppe et al., 2013). Pooling data across brains can be especially useful for iEEG/iEBS research which typically employs a very limited number of patients with idiosyncratic electrode placement.

iELVis currently supports five FreeSurfer based anatomical atlases, four cortical and one volumetric. Two of these, the Desikan-Killiany and Destrieux atlases (Desikan et al., 2006; Destrieux, Fischl, Dale, & Halgren, 2010), are based on the major gyri and sulci. The Desikan-Killiany atlas (Figure 4: A) divides the cortical surface into 35 areas, while the Destrieux atlas has a finer grained 75 area parcellation (Figure 4: B). These atlases are particularly useful for functional areas that largely follow gyrification patterns (e.g., primary motor and sensory cortex, Broca’s area). The other two cortical atlases are derived from resting state fMRI functional networks (Yeo et al., 2011). More specifically, Yeo and colleagues collected resting state fMRI data from 1000 neurotypical, young adults and grouped together cortical areas with high functional connectivity. They found two well-formed groupings of areas consisting of 7 and 17 networks (Figure 4: C-D). The 7-network atlas consists of the following groupings: default mode, frontoparietal, somatomotor, dorsal attention, ventral attention, limbic, and visual. These networks are somewhat broken up into the more fine-grained groupings of the 17-network atlas. For example, the 7-network atlas somatomotor network divides into dorsal and ventral somatomotor areas around the boundary of hand and face sensorimotor cortex.

**Figure 4:**
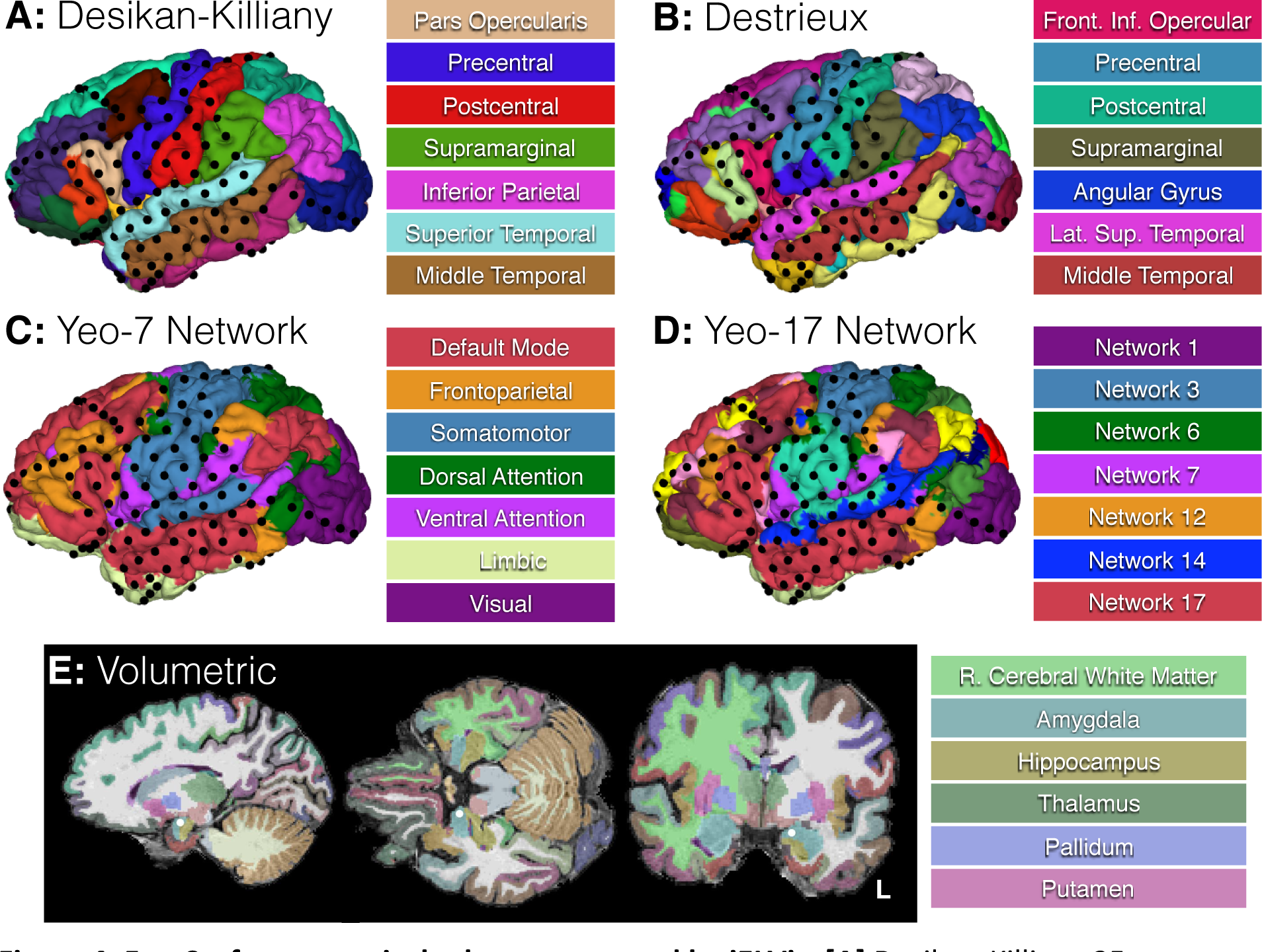
FreeSurfer anatomical atlases supported by iELVis. **[A]** Desikan-Killiany 35 area cortical atlas based on gyrification. **[B]** Destrieux 75 area cortical atlas based on gyrification. **[C]** Yeo 7 and **[D]** 17 area cortical atlas derived from resting state functional networks. **[E]** Volumetric atlas that includes subcortical as well as Desikan-Killiany cortical areas. Electrode locations are represented with black spheres (A-D) or white circles (E). Due to space limitations, the legends provide only a partial list of atlas areas. A full list of atlas parcellations is available on the iELVis wiki: http://episurg.pbworks.com/w/page/104391043/Mapping%20Electrodes%20to%20Atlases

The volumetric atlas labels 44 subcortical areas, including the amygdala, hippocampus, and white matter (Figure 4: E). In addition, cortical voxels are labelled according to the Desikan-Killiany atlas. As mentioned previously, depth electrode contacts that lie in cortical grey matter can be manually relabeled as subdural contacts. This will allow them to be localized using the higher resolution surface based atlases with iELVis.

## 5 MULTIMODAL OVERLAYS

Particularly in the context of invasive epilepsy monitoring, multiple modalities of brain mapping data are often acquired from the same individual. iELVis supports the simultaneous visualization of most such data. Specifically, it can visualize neuroimaging data (e.g., blood-oxygen-level dependent contrasts or cortical thickness), single contact electrode data (e.g. power changes), and bipolar contact electrode data (e.g. bipolar stimulation effects). Figure 5 displays an example of this. Specifically, it shows the overlay of three methods of mapping hand sensorimotor cortex: fMRI, iEEG, and iEBS. In addition, the iEEG and iEBS data are overlaid on the Yeo-17 area atlas to illustrate how well the Yeo et al. anatomically based functional networks compare with the patient’s functional mapping. All three functional mapping methods are in strong agreement (Figure 5: Columns A-B). The peak and boundaries of the fMRI statistical map are tightly consistent with that of the iEEG activations. The iEBS mapping is also highly consistent with the other two modalities, however it appears that iEBS includes an electrode that is a bit ventral and anterior of hand sensorimotor cortex because it was paired with another electrode placed squarely over hand sensorimotor cortex during stimulation.

**Figure 5:**
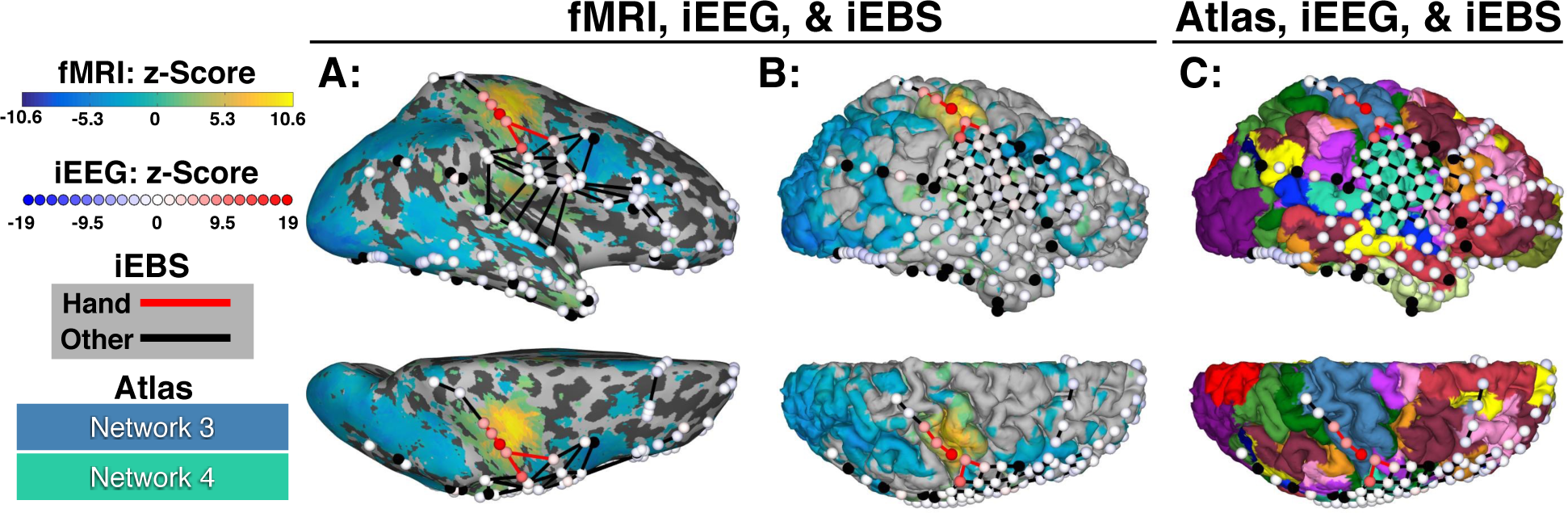
Using iELVis to visualize the localization of hand sensorimotor cortex using multiple modalities: fMRI, iEEG and iEBS. Maps are visualized on the inflated (Column A) and non-inflated (Columns B-C) pial surface. For the inflated pial surface, gyri and sulci are represented with light and dark grey, respectively. All data are shown in the same coordinate space (i.e., patient-specific FreeSurfer space, see Fig. 1). **[fMRI Data]** Parula (i.e., blue-green-yellow) color scale represents an fMRI statistical map with positive values indicating a greater blood-oxygen level dependent contrast during finger tapping versus rest. **[iEEG Data]** Blue-red color scale represents an iEEG-derived statistical map with positive values indicating greater high gamma band power during finger tapping versus rest. Spheres represent electrode locations. Black spheres indicate that an electrode provided no usable data. **[iEBS Data]** Red lines between electrodes indicate pairs of electrodes that produced hand movement or sensation when electrically stimulated. Black lines indicate that either no response or a non-hand sensorimotor response was elicited when the connected electrodes where stimulated. **[Yeo-17 Atlas]** Column C indicates the boundaries of various resting state functional networks according to FreeSurfer’s Yeo et al. 17 area atlas. The boundary between Networks 3 and 4 (i.e., the dorsal and ventral sensorimotor networks) is congruent with the fMRI, iEEG, and iEBS-defined ventral-most boundary of hand sensorimotor cortex.

Note that the Yeo-17 network dorsal somatomotor area captures the ventral boundary of hand somatomotor cortex (Figure 5: Column C). However, the atlas area extends far beyond the dorsal boundary of the hand region and likely includes other somatomotor regions (e.g., arm and leg).

Finally, note that the visualization of electrode data on the inflated pial surface (Figure 5: Column A) makes it possible to relate electrode data to sulcal neuroimaging data and, potentially, to visualize penetrating depth electrodes that lie in sulcal grey matter. This is useful not only because most of the cortical surface is sulcal (Valiante, 2012) and often obscured in standard iEEG visualizations, but also because the inflated surface provides a better sense of the surface-based distance (as opposed to Euclidean) between electrodes. This could be very helpful for understanding the spread of cortical activity such as seizures (Jenssen, et al., 2011; Schevon et al., 2012) or how functional interactions between cortical areas varies as a function of distance (Keller, Bickel et al., 2011).

## 6 DISCUSSION

Despite the increasing importance and popularity of iEEG and iEBS research, public software tools for dealing with the unique technical challenges of this work are limited. With the iELVis toolbox, we have addressed the need for an open source, well-documented software package that solves many common iEEG/iEBS technical challenges. iELVis provides a family of MATLAB functions for identifying the anatomical locations of electrodes with mm-scale resolution, for surface-based mapping of electrode locations to an average brain for group analyses, for mapping electrode locations to five anatomical atlases, and for simultaneously, interactively visualizing electrode and neuroimaging data. The toolbox works on all major operating systems, is well-documented, includes well-developed tutorial and documentation materials, and is easy to integrate with other popular neuroscience freeware packages such as EEGLAB (Delorme & Makeig, 2004; Delorme et al., 2011), FieldTrip (Oostenveld, Fries, Maris, & Schoffelen, 2011), and SPM (http://www.fil.ion.ucl.ac.uk/spm/).

While the iELVis toolbox meets the typical, basic localization needs of iEEG/iEBS research, the current version of the toolbox has some shortcomings. In particular, an issue with the surface based mapping to the average brain is that small differences in electrode locations in the patient’s native space can result in large differences in average brain coordinates if an electrode lies near gyral borders (e.g., near the boundary between the temporal and frontal lobes). Similarly, small differences in electrode locations can affect the anatomical region assigned to electrodes that lie near the borders of different atlas regions. A simplistic way to deal with this would be to remove these boundary electrodes from group analyses since their anatomical location is uncertain. Alternatively, subdural electrode data may be spatially smeared across nearby cortical vertices to account for the uncertainty of electrode locations (Kadipasaoglu et al., 2014; Kubanek & Schalk, 2015). A complication with this is approach is that it is unclear what the smearing kernel should be and such smearing fails to deal with volume conduction. The best alternative may be to localize the neural sources of the electrode data (Akalin Acar, et al., 2011) and to map those data to the average brain or atlases. This method could remove the need for a somewhat arbitrary smoothing kernel and would account for volume conduction. In particular, it could identify sulcal activity that likely contributes to subdural iEEG/iEBS data (Towle, Carder, Khorasani, & Lindberg, 1999).

An additional shortcoming of iELVis is that it does not implement any method for correcting depth electrodes for postimplant brain shift when the depth electrodes are implanted along with subdural electrodes that require a craniotomy. As already mentioned, the degree of brain shift near depth contacts is typically minor given that they are anchored to the skull and are usually distant from the craniotomies that produce the most significant brain shift. However, it would be useful to have tools for depth electrode brain-shift correction as sometimes depth electrodes lie near the craniotomy and if brain shift is severe it may affect even deep structures. A more significant limitation is that the automatic FreeSurfer pial surface construction, which iELVis depends on, often fails near abnormal brain regions (e.g., tumors or previous resections) and typically underestimates the medial extent of the medial temporal pial surface. The pial surface can potentially be manually corrected in FreeSurfer, however it may be that alternative surface based electrode localization algorithms (e.g., Hermes, Miller, Noordmans, Vansteensel, & Ramsey, 2010) might be more effective in cases of significant failure. Finally, an inconvenience of iELVis is its current dependence on multiple non-MATLAB based third-party software packages. This complicates installation and maintenance of iELVis. However, in our experience these dependencies require minimal additional effort, especially since many laboratories already use FreeSurfer and FSL for neuroimaging analysis, and the core functionalities of these decade-old packages are quite stable.

In the future, we plan to develop iELVis further to address some of these limitations and to add new functionality. For example, we aim to add tools to quantify the degree of confidence that an electrode can be assigned to a particular anatomical region in order to identify electrodes whose anatomical locus is in doubt (Mercier et al., 2016). Moreover, we hope to make it possible to spatially smooth electrode data over the cortical surface or to localize the neural generators of iEEG data in order to improve group analyses. Finally, we hope to add compatibility with additional brain atlases such as probabilistic maps of visual areas (L. Wang, Mruczek, Arcaro, & Kastner, 2015b) and individual-data derived resting-state fMRI network parcellations (Fox et al., 2016; D. Wang et al., 2015a), and to remove some dependencies on third party software. We also welcome contributions from other iEEG/iEBS researchers and have established a users group and contribution guidelines to facilitate this (http://episurg.pbworks.com).

## ACKNOWLEDGMENTS

We thank the patients for consenting to provide the data that made this toolbox possible. Moreover, we thank Miklos Argyelan, Kathrin Müsch, Taufik Valiante, and Jaime Gomez for helping to debug and to develop this software. This work was supported by the Swiss National Science Foundation (grants PBGEP3_139829 and P300P3_148388 to PM), the Page and Otto Marx Jr. Foundation (to ADM), and by the Natural Sciences and Engineering Research Council of Canada (RGPIN-2014-04465 to CJH).

## Footnotes

1 In previous papers (Dykstra et al., 2011, Yang, Wang, et al., 2012) the smoothed pial surface was called the “dural surface.” We think “leptomeningeal surface” is more accurate given that subdural electrodes lie below the dural membrane.

